# A low-input high resolution sequential chromatin immunoprecipitation method captures genome-wide dynamics of bivalent chromatin

**DOI:** 10.1101/2023.09.18.558170

**Authors:** William Ho, Janith A. Seneviratne, Eleanor Glancy, Melanie A. Eckersley-Maslin

## Abstract

**Background:** Bivalent chromatin is an exemplar of epigenetic plasticity. This co-occurrence of active-associated H3K4me3 and inactive-associated H3K27me3 histone modifications on opposite tails of the same nucleosome occurs predominantly at promoters where it poises them for future transcriptional upregulation or terminal silencing. We know little of the dynamics, resolution, and regulation of this chromatin state outside of embryonic stem cells where it was first described. This is partly due to the technical challenges distinguishing bone-fide bivalent chromatin, where both marks are on the same nucleosome, from allelic or sample heterogeneity where there is a mix of H3K4me3-only and H3K27me3-only mononucleosomes.

**Results:** Here, we present a robust and sensitive method to accurately map genome-wide bivalent chromatin along with all necessary controls from as little as 2 million cells. We optimised and refined the sequential ChIP protocol which uses two sequential overnight immunoprecipitation reactions to robustly purify nucleosomes that are truly bivalent and contain both H3K4me3 and H3K27me3 modifications. Our method generates high quality genome-wide maps with strong peak enrichment and low background which can be analysed using standard bioinformatic packages. Using this method, we detect twice as many bivalent regions in mouse embryonic stem cells as previously identified, bringing the total number of bivalently marked gene promoters to 8,373. Furthermore, profiling Dppa2/4 knockout mouse embryonic stem cells which lose both H3K4me3 and H3K27me3 at approximately 10% of bivalent promoters, demonstrated the ability of our method to capture bivalent chromatin dynamics.

**Conclusions:** Our optimised sequential reChIP method enables high-resolution genome-wide assessment of bivalent chromatin together with all required controls in as little as 2 million cells. We share a detailed protocol and guidelines that will enable bivalent chromatin landscapes to be generated in a range of cellular contexts, greatly enhancing our understanding of bivalent chromatin and epigenetic plasticity beyond embryonic stem cells.

## Background

The chromatin landscape of cells not only shapes cellular identity but also enables how cells are able to respond and adapt to a changing environment. Amongst the multitude of different layers of organisation, histone post-translational modifications are tightly associated with the activity and accessibility of the underlying DNA sequence. In particular, tri-methylation of lysine 4 on histone 3 (H3K4me3) is tightly correlated with active promoters, whilst tri-methylation of lysine 27 of histone 3 (H3K27me3) is associated with heterochromatin and gene repression. Remarkably these two seemingly opposing histone modifications can be found on opposite tails of the same nucleosome where it is thought to keep the underlying DNA sequence poised and amenable to future activation or repression (reviewed in (1,2)). In mouse embryonic stem cells, removing bivalent chromatin results in the accumulation of tightly repressive DNA methylation and the inability of the genes to be activated in a timely manner upon differentiation (3–7). Therefore, bivalent chromatin is a classic exemplar of molecular plasticity, by priming genes for the future and facilitating cell adaptation. However, bivalent chromatin has been predominantly studied in the context of mouse embryonic stem cells (ESC) where it was first described (8,9). Consequently, our current understanding of the distribution and dynamics in other cell types and species remains limited. This is partly due to technical challenges associated with accurately detecting this important structure.

A major challenge in studying bivalent chromatin is that the co-occurrence of active H3K4me3 and inactive H3K27me3 histone modifications needs to be distinguished from instances where the histone modifications occur on different alleles in the cell or in different cells within a mixed population (Figure 1A). Consequently, performing independent chromatin immunoprecipitation (ChIP) or CUT&RUN-related methods separately for H3K4me3 and H3K27me3 and then overlapping peaks *in silico* is not sufficient to be absolutely certain the region is indeed bivalent and not a consequence of allelic or cellular heterogeneity. This becomes even more of a challenge when analysing complex systems such as developing tissues or patient cancer samples. Previous studies in human T cells have suggested that as many as 14% of bivalent regions called using independent ChIPs are false-positives (10). To address this, sequential ChIP or ChIP-reChIP approaches have been developed (10–15), whereby the chromatin purified from a first immunoprecipitation reaction (e.g. H3K4me3) is used as input into a second immunoprecipitation reaction for a different modification (e.g. H3K27me3). Theoretically, only chromatin with both marks of interest are purified in this way. However, these protocols typically require tens of millions of cells as input and so are not always feasible. Studies often perform the reChIP in just one direction (e.g. H3K4me3 followed by H3K27me3 but not *vice-versa*) which has important consequences as the resulting datasets frequently suffer from “signal carry-over” from the first ChIP into the second, leading to many false-positives. Moreover, poor signal-to-noise makes data interpretation and downstream analysis complex. Recently, multi-tagmentation methods have been described to simultaneously map multiple histone modifications in single-cells (16–18), yet these methods required custom reagents such as different barcoded Tn5 complexes or nanobodies, and complex data-analysis pipelines. Therefore, there is a need for sensitive, robust and cost-effective methods to accurately detect bivalent chromatin in low cell numbers that can use existing standardised downstream data-analysis approaches.

**Figure 1:**
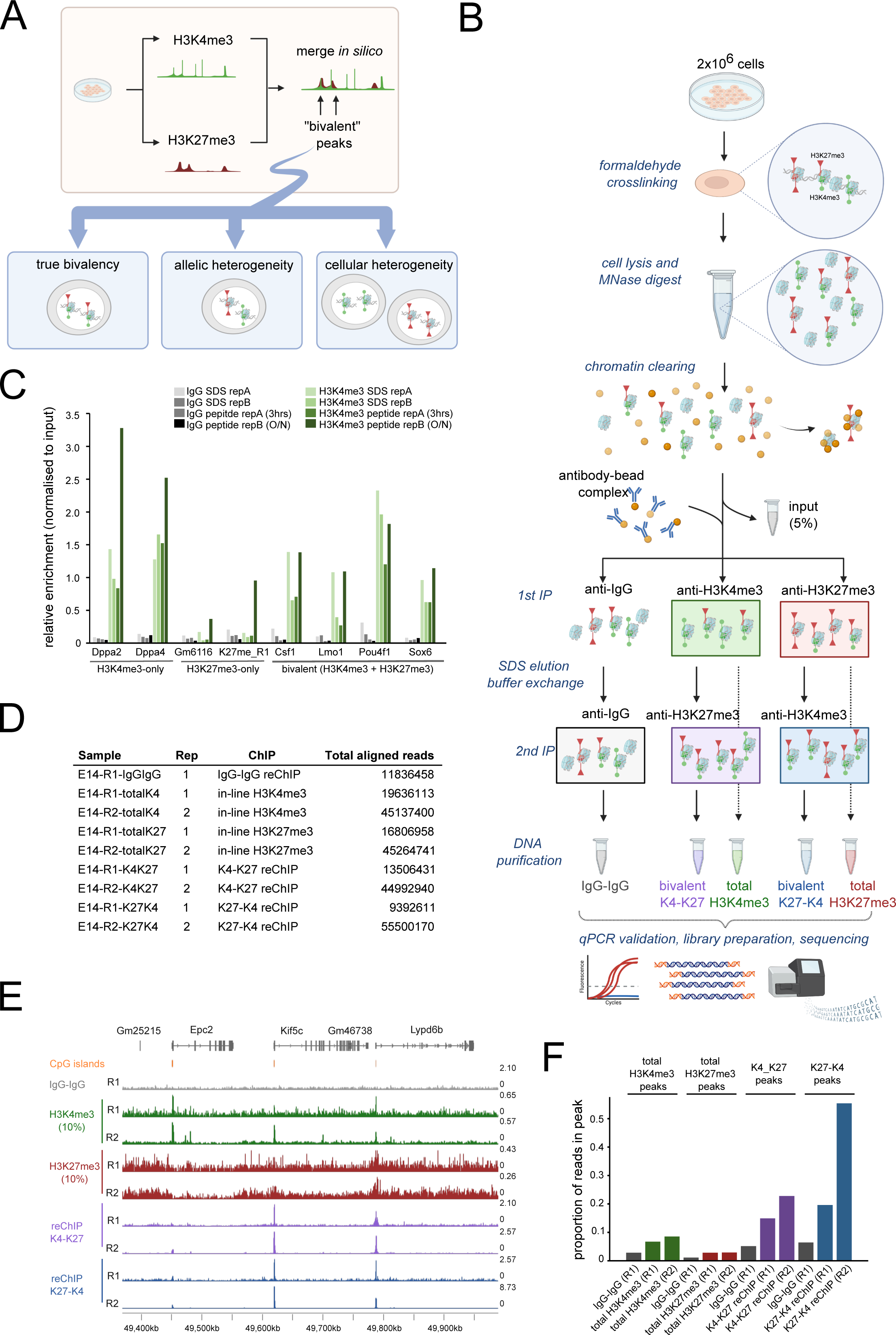
Development of an optimised ChIP-reChIP protocol to robustly measure bivalent chromatin. (A) Potential limitations in using independent total H3K4me3 (green, circles) and total H3K27me3 (red, triangles) datasets in distinguishing bone-fide bivalent chromatin, where the two marks occur on the same nucleosome, from allelic and cellular heterogeneity. (B) overview of sequential ChIPreChIP protocol. (C) Single H3K4me3 (green) and IgG control (grey) ChIP-qPCR analysis comparing SDS-based elution (light) from peptide elution (dark). Two H3K4me3-only (Dppa2, Dppa4), two H3K27me3-only (Gm6116, K27me_R1) and four bivalently marked loci (Csf1, Lmo1, Pou4f1, Sox6) were analysed. Enrichment values normalised to input are shown. (D) Summary table of samples sequenced in E14 mouse embryonic stem cells indicating replicate, ChIP type and total aligned reads (E) Genome browser view of reChIP datasets including IgG-IgG reChIP control (grey), in-line total H3K4me3 (green, row 2 and 3), in-line total H3K27me3 (red, row 4 and 5) and bivalent reChIP for H3K4me3 followed by H3K27me3 (K4-K27, purple, row 6 and 7) or vice-versa (K27-K4, blue, row 8 and 9). Two biological duplicates (R1 and R1) are shown for all but IgG-IgG libraries. CpG islands are denoted by orange bars. (F) FRiP scores showing proportion of reads within peaks for each individual sample. IgG-IgG (grey) is shown to get background levels.

Here, we present a highly-optimised sequential ChIP (reChIP) methodology for accurately detecting bivalent chromatin along with all required controls from just 2 million cells. From one sample, our refined method generates 5 datasets including the 2 reChIP datasets (H3K4me3 is followed by H3K27me3 and *vice versa*) and 3 control datasets (IgG-IgG background reChIP, in-line total H3K4me3 and in-line total H3K27me3). By applying our method in mouse ESCs we detected 7,714 high confidence bivalent chromatin regions which occurred predominantly at CpG-rich promoters. Importantly, in addition to 97% of previously annotated bivalent genes (14), our method revealed an additional 4,780 bivalent gene promoters, more than doubling the catalogue of mouse ESC bivalent genes. Lastly, we validated the sensitivity of our method by profiling ESCs lacking the epigenetic priming factors *Dppa2* and *Dppa4* which are required for maintaining bivalent chromatin at a subset of promoters (3,4). This confirmed the ability of our method to detect dynamic changes in bivalent chromatin. In summary our method provides a much-needed resource for researchers wishing to accurately map bivalent chromatin landscapes from as little as 2 million cells.

## Results

### Development of an optimised ChIP-reChIP protocol to robustly measure bivalent chromatin

A challenge in studying bivalent chromatin is that aligning independently generated single H3K4me3 and H3K27me3 datasets *in silico* is theoretically insufficient to distinguish true bivalency (where both marks are present on the same chromatin fragment) from allelic or cellular heterogeneity (where marks are present on different alleles or in different cells within the population) (Figure 1A). To address this, reChIP (also known as sequential ChIP or ChIP-reChIP) approaches have been used (10–15,19,20) whereby following the first immunoprecipitation the eluted sample is sequentially immunoprecipitated with a different antibody against the second mark of interest. However, many limitations exist with current protocols. Unfortunately, many existing reChIP protocols require large (>10 million cells) amount of starting material per reaction (10,11,14,19,20), often the experiments are performed in just one of the two directions (20). This is a major issue as many false positives can confound the results due to “signal carry-over” whereby enrichment from the first ChIP carries through into the second ChIP. Moreover, variable data quality with low signal to noise has traditionally made downstream bioinformatic analysis of bivalent regions challenging. In order to address these points, we optimised the reChIP protocol to give a high signal to noise ratio with both qPCR and high-throughput sequencing readouts from just 2 million cells (Figure 1B). Critically, we advise the reChIP be carried out in both orientations (H3K4me3 followed by H3K27me3 and *vice versa*). The method was optimised using serum/LIF cultured E14 mouse Embryonic Stem Cells (ESCs) given their well-defined distribution of bivalent chromatin (8,9,14,15). Importantly our method produces high quality data that can be analysed with commonly-used bioinformatic tools. A full detailed protocol accompanies this paper (Supplemental File 1).

The 3-day workflow is shown in Figure 1B. Briefly, cells are treated with formaldehyde to cross-link chromatin and 2 million cell aliquots stored at −80°C for up to 6 months. Cells are gently lysed and treated with MNAse to generate predominantly mononucleosomes and chromatin pre-cleared by incubating with pre-washed dynabeads for 3 hours at 4°C to reduce non-specific binding. During this time, the antibody-dynabead complexes are formed for the IgG control, H3K4me3 and H3K27me3 immunoprecipitations. 5% of the precleared chromatin is set aside as an input control and the remainder split across the three tubes of antibody-dynabead complexes for the first overnight immunoprecipitation at 4°C.

Following the first immunoprecipitation, the chromatin-antibody-dynabead complexes are thoroughly washed to remove any non-specific binding prior to chromatin elution. Traditionally, reChIP protocols typically use one of two approaches to elute chromatin from beads. DTT- or SDS-based elution buffers function by dissociating the affinity interactions upon which the immunoprecipitation is based but requires additional dilution and/or cleanup steps to ensure compatibility with a second immunoprecipitation reaction (11–14,19,20). An alternative is to use high concentration of modified histone tail peptides to compete with antibody binding sites (10). We compared these two approaches to elute chromatin in single H3K4me3 ChIPs. SDS elution performed well in terms of specificity and signal to noise ratio. While the 3-hour peptide competition gave similar results to SDS, the amount of unspecific background signal increased when incubated overnight (Figure 1C). After considering costs and availability of commercial peptides, we decided to implement SDS elution in our final protocol. To mitigate against the presence of SDS inhibiting subsequent antibody binding events, we diluted the chromatin and performed a buffer exchange using 3 kDa molecular weight filters. From the first immunoprecipitation reaction, 10% of the sample representing in-line total H3K4me3 or total H3K27me3 control is set aside. The second immunoprecipitation is then performed overnight using the alternate antibody so that the reChIP is performed in both directions: H3K4me3 followed by H3K27me3 (K4-K27) and H3K27me3 followed by H3K4me3 (K27-K4). As a negative control, IgG followed by IgG (IgG-IgG) is also performed to control for non-specific enrichment during the reChIP assay. Chromatin is then eluted in SDS-elution buffer, formaldehyde crosslinks reversed, RNA and proteins degraded, and enriched DNA fragments purified ready to be processed for qPCR analysis and/or high throughput sequencing.

### Identification of 7,714 high confidence bivalent regions in mouse embryonic stem cells

To date bivalent chromatin is best understood in mouse ESCs. Therefore, we used this model to test our refined method. In total 9 datasets were generated from two biological replicates (Figure 1D). These included two in-line H3K4me3 single ChIPs, two in-line H3K27me3 single ChIPs, two each of K4-K27 and K27-K4 reChIPs, and one IgG-IgG replicate, with replicate 2 sequenced at a higher depth than replicate 1.

Initial data inspection of our reChIP datasets revealed strong peak distribution of reads for the K4-K27 and K27-K4 reChIP samples at known bivalent regions with low intervening background signal (Figure 1E). Furthermore, peaks were observed in the in-line total H3K4me3 and total H3K27me3 samples, albeit these signals were noisier. This is not likely due to sequencing coverage, which for replicate 1 was over 45 million aligned reads (Figure 1D), but rather due to the lower starting material for library preparation of these samples which only correspond to approximately 60,000 cells. We had sequenced the two biological replicates at different depths ranging from approximately 10 million through to 55 million aligned reads (Figure 1D). From the higher coverage replicate, we performed *in silico* downsampling analysis from which we concluded that 15-20 million reads was a good compromise between number of peaks detected versus sequencing cost, and that sequencing beyond this predominantly split peaks into multiple smaller peaks and/or called non-convincing peaks (data not shown). Supporting this, even with approximately 10 million mapped reads there were clear peaks in the reChIP samples (Figure 1E). To get a measure of the specificity of our assay, we calculated the fraction of reads in called peak regions (FRiP score) which is commonly used in ATAC-seq analysis to determine library quality. Notably, all reChIP samples had very high FRiP scores, while IgG-IgG scores were all less than 0.1 (Figure 1F). This indicates a very high and specific enrichment and low background of these reChIP libraries.

We called peaks separately for K4-K27 and K27-K4 reChIPs (see methods) obtaining 25,540 and 36,235 peaks respectively of which 21,857 peaks were shared (Figure 2A). We subsequently classified these peaks based on whether they were shared with total H3K4me3 and/or total H3K27me3 datasets. Bivalent peaks were classified into four categories: high-confidence peaks also overlapped peaks in both total H3K4me3 and H3K27me3 datasets; K4-biased and K27-biased overlapped peaks only in total H3K4me3 or H3K27me3 respectively; and low-confidence did not share a peak in either total H3K4me3 nor H3K27me3 (Supplemental Figure 1A-E). When we stratify bivalent peaks with these criteria using our in-line total H3K4me3 and H3K27me3 single ChIPs, half of the peaks (n=11,334 of 21,857) were classified as K4-biased with another 2,774 classified as high-confidence (Figure 2B, left). Since the “in-line” total ChIPs represent approximately 60,000 cells, we also stratified the 21,857 reChIP peaks using independent total ChIPs from approximately 10 million cells (Figure 2B, right) we previously generated using the same cell line (4). When using independently derived total ChIP-seq datasets, the number of high-confidence bivalent peaks increased three-fold to 7,714. Of note, the majority of these were due to re-classification of peaks categorised as K4-biased using the in-line total ChIPs. This suggests that the low input H3K27me3 single ChIP-seq dataset was masking many high-confidence bivalent peaks. Importantly, almost all of the 2,774 high-confidence bivalent peaks called using the in-line total ChIPs were contained within the 7,714 high-confidence bivalent peaks called using independent total ChIPs, demonstrating the increased sensitivity of our assay and very low false-positive rate. However, many high confidence bivalent regions are missed when using the in-line total ChIPs, likely due to the increased signal-to-noise ratio in these datasets. This is particularly important for the H3K27me3 single ChIP due to it’s broader distribution and consequently more dispersed signal which is challenging to capture in low-input protocols. Thus, while using the in-line total H3K4me3 and total H3K27me3 controls is suitable for accurately detecting some high-confidence bivalent regions, use of independent total-ChIP datasets facilitates high-confidence classification of approximately three times as many peaks. Therefore, when possible, we recommend generating independent total H3K4me3 and total H3K27me3 datasets to capture as many high confidence bivalent regions as possible.

**Figure 2:**
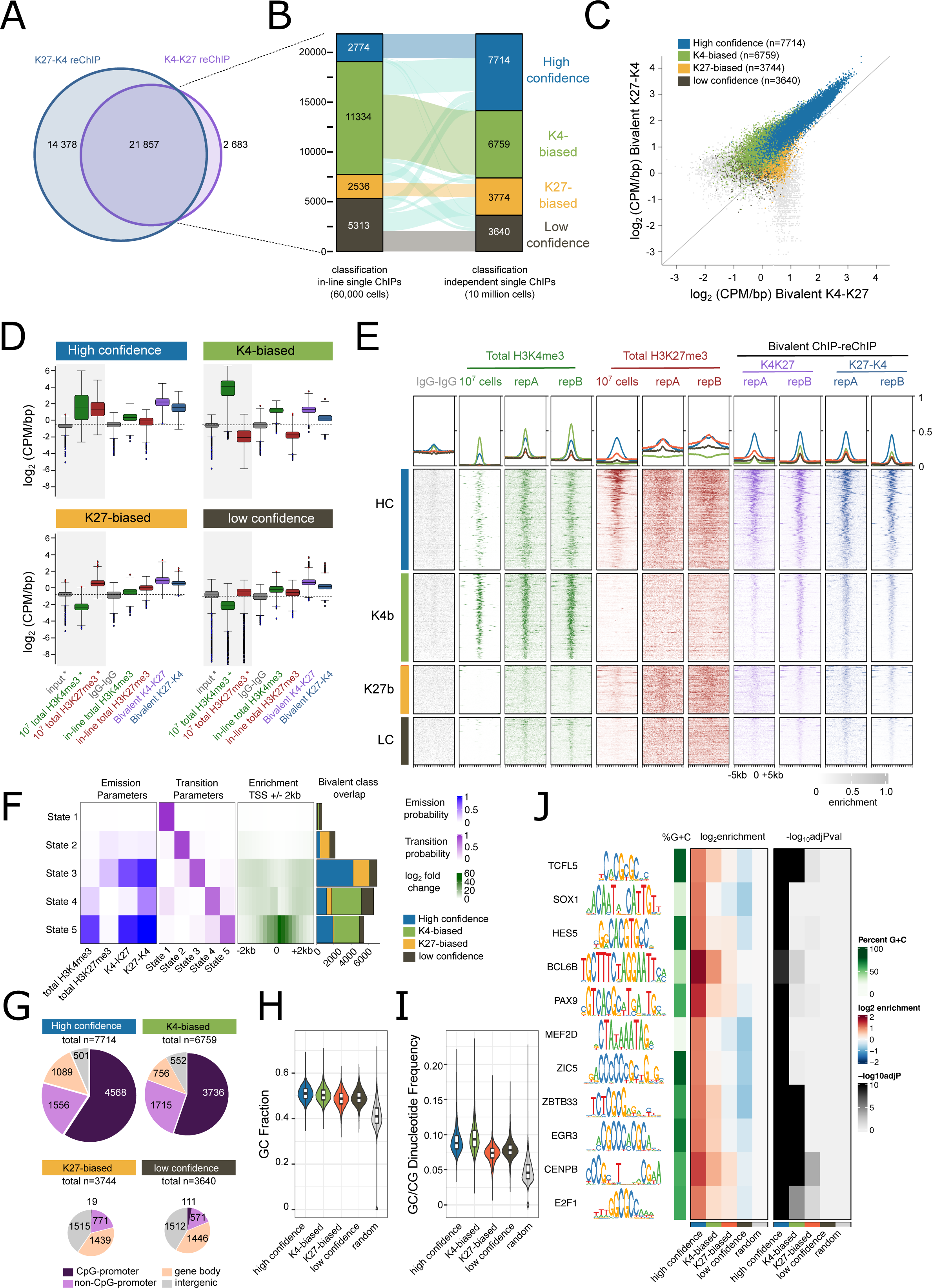
Identification of 7,714 high confidence bivalent regions in mouse embryonic stem cells. (A) Overlap between K27-K4 (blue) and K4-K27 (purple) reChIP datasets. (B) Alluvial plot showing classification of 21,857 peaks that overlap in both K27-K4 and K4-K27 reChIP datasets using in-line total H3K4me3 and H3K27me3 ChIPs from approximately 60,000 cells (left) or separate total H3K4me3 and H3K27me3 single ChIPs from approximately 1 million cells (right) from GSE135841 (4). Peaks were classified as high confidence (overlap peak in both total H3K4me3 and total H3K27me3, blue), K4-biased (overlap peak in only total H3K4me3, green), K27-biased (overlap peak in only H3K27me3, orange) or low confidence (does not overlap peak in either H3K4me3 or H3K27me3, brown). (C) Scatter plot showing log_2_CPM/bp values for bivalent K4-K27 (x-axis) and bivalent K27-K4 (y-axis) datasets for all bivalent peaks highlighting high confidence (blue), K4-biased (green), K27-biased (orange) and low confidence (brown) peaks. (D) Box-whisker plots showing log_2_CPM/bp values for high confidence (top left), K4-biased (top right), K27-biased (bottom left) and low confidence (bottom right) peaks in independent total H3K4me3 and total H3K27me3 datasets from GSE135841 (4) (denoted by * and shaded grey background) or the in-line total and reChIP datasets generated in this study. (E) Enrichment heatmaps showing CPM/bp normalised read densities for high confidence (top row), K4-biased (second row), K27-biased (third row) and low confidence (bottom row) peaks after scaling for all datasets analysed. Peaks were extended by 5kb upstream and downstream. Values surpassing the 99^th^ percentile have been masked for visualisation. 10^7^ samples refers to independent total H3K4me3 and total H3K27me3 datasets from GSE135841 (4) (F) 5-state chromHMM models using pooled replicates for in-line total H3K4me3, in-line total H3K27me3 and K4-K27 and K27-K4 reChIP datasets showing emission (left) and transmission (second from left) parameters, enrichment across TSS +/-2kb and overlap with high confidence (blue), K4-biased (green), K27-biased (orange) and low-confidence (brown) bivalent regions (right). (G) Genomic features associated with the four classes of bivalent regions. (H, I) Violin plots showing GC fraction (H) and GC-CG dinucleotide frequency (I) within regions compared to random subset of genomic regions with same number as high confidence regions. All comparisons are statistically significant after multiple testing (Benjamini-Hochberg correction). (J) Motif enrichment for the four classes of bivalent peaks compared to random genomic sequences. Those with log_2_enrichment over random sequences >1 are shown, along with their enrichment scores and −log_10_Adjusted P-value.

Overlapping peaks *in silico* is often used as a proxy to define bivalent chromatin (Figure 1A). To assess the fidelity of this approach in accurately calling bivalent regions, we overlapped total H3K4me3 and total H3K27me3 peaks (called using the independent 10 million cell dataset) *in silico* to obtain 7,868 overlapping regions. Of these, 7,613 (96.8%) and 7,801 (99.1%) regions also overlapped the bivalent K4K27 and K27K4 reChIP peaks respectively, and 7,611 (96.7%) overlapped a peak in both reChIP datasets (Supplemental Figure 1F). While this overlap is very strong, the increased sensitivity of our reChIP assay is evident through the detection of an additional 13,632 peaks in both K4-K27 and K27-K4 directions. This data implies that, at least in mouse embryonic stem cells, using independently derived total ChIP-seq datasets is sufficient to detect bivalent regions with a very low false-positive rate but that its sensitivity is limited. It remains unknown if this holds true in other cell types or complex tissues.

The 7,714 high confidence bivalent regions classified using independent total H3K4me3 and total H3K27me3 had the highest enrichment in both K4-K27 and K27-K4 reChIP orientations (Figure 2C, D). The reChIP signal at K4-biased, K27-biased and low-confidence bivalent regions was lower (Figure 2C, D). Unlike the broad total H3K27me3 peaks, bivalent reChIP peaks were sharp and narrow (Figure 2E). This demonstrates the specificity of our approach in enriching for chromatin fragments containing both modifications of interest and the absence of carry-over of the broader H3K27me3 signal particularly in the K27-K4 reChIP dataset. Orthogonal unbiased Hidden Markov Model approaches (21) using in-line totals and reChIPs identified chromatin states that matched our peak-centric classifications (Figure 2F). From the 5-state chromatin model we observed that state 3 was enriched for our high confidence bivalent promoters, whilst state 4 and 5 represented a mix of high confidence and K4-biased bivalent regions occurring around and at gene promoters respectively. This analysis confirms the validity of these bivalent peak subclasses (Figure 2F).

Next, we wanted to determine the distribution of bivalent domains across different genomic elements. The high confidence and K4-biased bivalent regions had the highest levels of reChIP enrichment (Figure 2C, D) and were predominantly located at CpG (59.2% and 55.3% respectively) and non-CpG promoters (20.1% and 25.4% respectively) (Figure 2G), consistent with previous studies (1,2,8). In contrast, K27-biased and low confidence bivalent regions had lower levels of enrichment (Figure 2C, D) and occurred predominantly at gene bodies and intergenic regions (Figure 2G). Sequence analysis revealed that all classes of bivalent regions had a higher GC content (Figure 2H) and CpG frequency (Figure 2I) than a size-matched random set of genomic regions. Motif analysis revealed a strong enrichment for motifs associated with developmental regulation including SOX1, HES5, PAX9 and ZIC5 (Figure 2J) in the high confidence and K4-biased but not the K27-biased and low confidence bivalent regions. In summary our method is able to robustly detect thousands of high-confidence bivalent regions in mouse embryonic stem cells occurring predominantly at CpG-promoters enriched for developmental transcription factor binding sites.

### Catalogue of 8,373 high-confidence bivalent genes in mouse embryonic stem cells

Given the strong overlap between high confidence bivalent regions and gene promoters, we next analysed gene promoters specifically. Using the independent 10 million cell totals for classification revealed 8,373 gene promoters that overlapped a HC-bivalent peak. Importantly this included 3,593 of 3,699 (97%) previously annotated bivalent genes in mouse embryonic stem cells (14) (Figure 3A). Of note, however, is our improved sensitivity in detecting an additional 4,780 high-confidence bivalent gene promoters. Representative H3K4me3-only, high-confidence, K4-biased, K27-biased and low-confidence bivalent promoters are shown in Supplemental Figure 1A-E.

**Figure 3:**
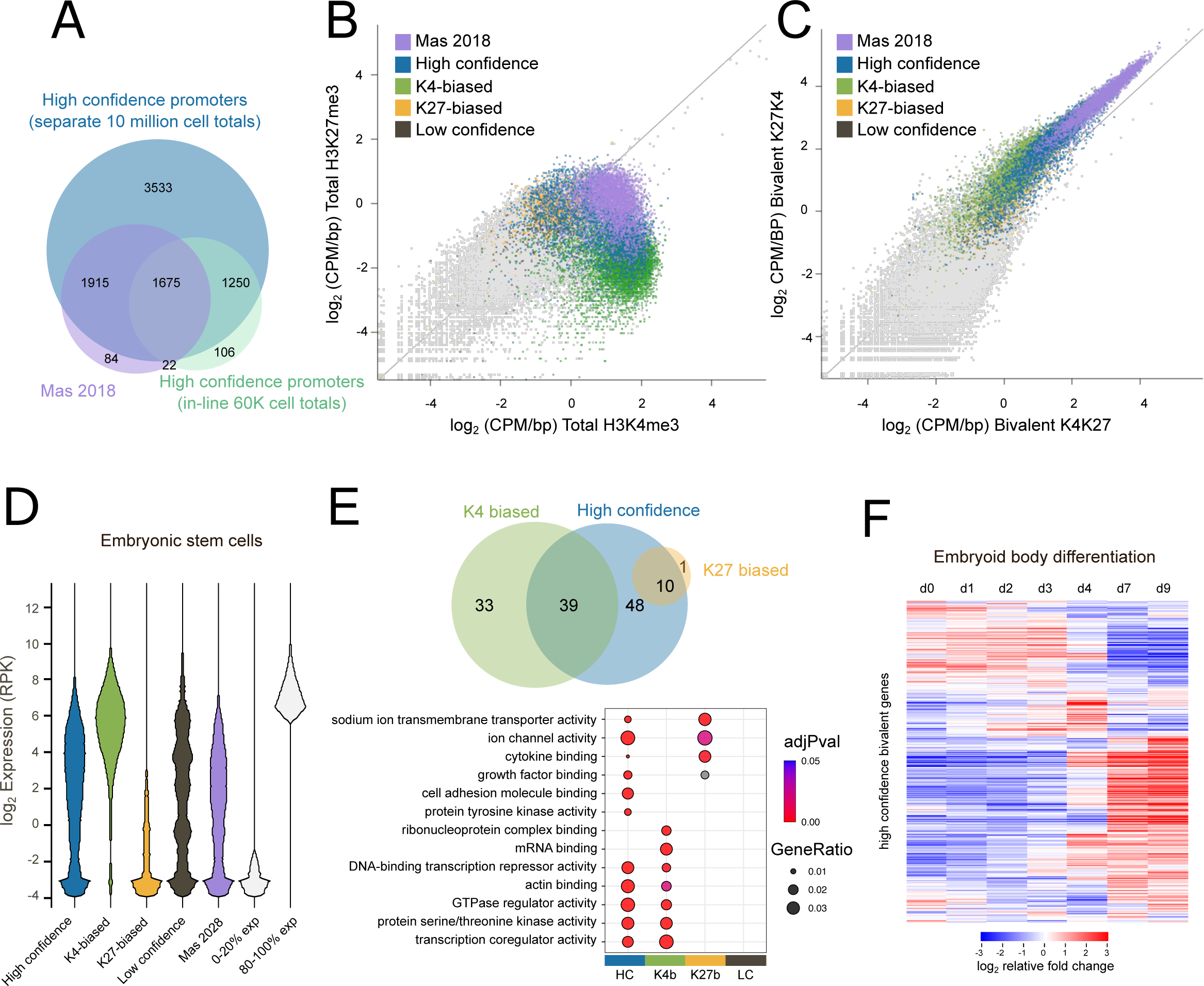
Catalogue of 8,383 bivalent genes in mouse embryonic stem cells. (A) Overlap of promoters classified as high-confidence bivalent in this study using in-line 60K total H3K4me3 and total H3K27me3 (aqua), independent 10 million cell total H3K4me3 and total H3K27me3 (GSE135841) (4) (blue) or previously published bivalent gene set (Mas et al. 2018) (14). Full list of bivalent promoter classifications are available in Supplemental Table 3. (B,C) scatterplot showing log_2_enrichment (CPM/bp) of (B) in-line total H3K4me3 (x-axis) and in-line total H3K27me3 (y-axis) or (C) bivalent K4-K27 (x-axis) and K27-K4 (y-axis) reChIP datasets for all promoters highlighting those that overlap different classes of bivalent peaks defined using 10 million cell total H3K4me3 and total H3K27me3. (D) log_2_ gene expression levels in mouse embryonic stem cells for four different classes of bivalent genes and previously annotated bivalent genes (14). Expression of the bottom 20% and top 20% are shown as a comparison. Gene expression data reanalysed from GSE135841. (E) Gene Ontology analysis showing overlap of enriched terms in the four classes of bivalent genes (top) and gene ratios and adjusted P-value of selected terms (bottom). The full list of enriched terms is available in Supplemental Table 4. (F) log_2_ fold change in gene expression levels for high confidence bivalent genes across 9 days of embryoid body differentiation. Each gene has been normalised separately across the time series. Gene expression data reanalysed from (GSE135841).

The high confidence associated bivalent gene promoters had the highest levels of total H3K4me3 and total H3K27me3 (Figure 3B) and bivalent K4-K27 and K27-K4 reChIP enrichment (Figure 3C). The high confidence bivalent genes were expressed at low yet detectable levels in pluripotent mouse embryonic stem cells (Figure 3D). In contrast the K4-biased bivalent genes had higher expression values, consistent with their enrichment for total H3K4me3 but not total H3K27me3, while expression of K27-biased bivalent genes was barely detectable (Figure 3D). The high confidence bivalent genes were enriched in pathways relating to ion channel activity, growth factor binding, cell adhesion, transcriptional regulation and protein kinase activity (Figure 3E) of which many were shared with the K4-biased or K27-biased classes (but not both). In line with current models (1,2,22), high confidence bivalent genes were dynamically expressed upon differentiation resolving to either an active or repressed state (Figure 3F). Therefore, our data supports the current model of bivalent chromatin marking developmental genes in embryonic stem cells for future activation or repression.

### Profiling bivalent chromatin dynamics in DPPA2/4 knockout mouse embryonic stem cells

Lastly, we confirmed the sensitivity of our method to detect changes in bivalent chromatin by profiling mouse embryonic stem cells deficient for the epigenetic priming factors DPPA2 and DPPA4 (4). We and others recently reported that Dppa2/4 are required to maintain bivalent chromatin at a subset of bivalent genes (3,4) (Figure 4A). To test the dynamic sensitivity of our method, we profiled two wild-type (WT) and two Dppa2/4 double knockout (DKO) clones using our refined method. Our reChIP datasets recapitulated previous observations where total H3K4me3 and total H3K27me3 signals were lost at Dppa2/4-dependent bivalent genes yet retained at Dppa2/4-independent genes in the Dppa2/4 knockout cells (Figure 4B). Importantly, this was also observed in the both bivalent K4-K27 and K27-K4 reChIP directions. This highlights the ability of our improved method to detect dynamics of bivalent chromatin between different samples.

**Figure 4:**
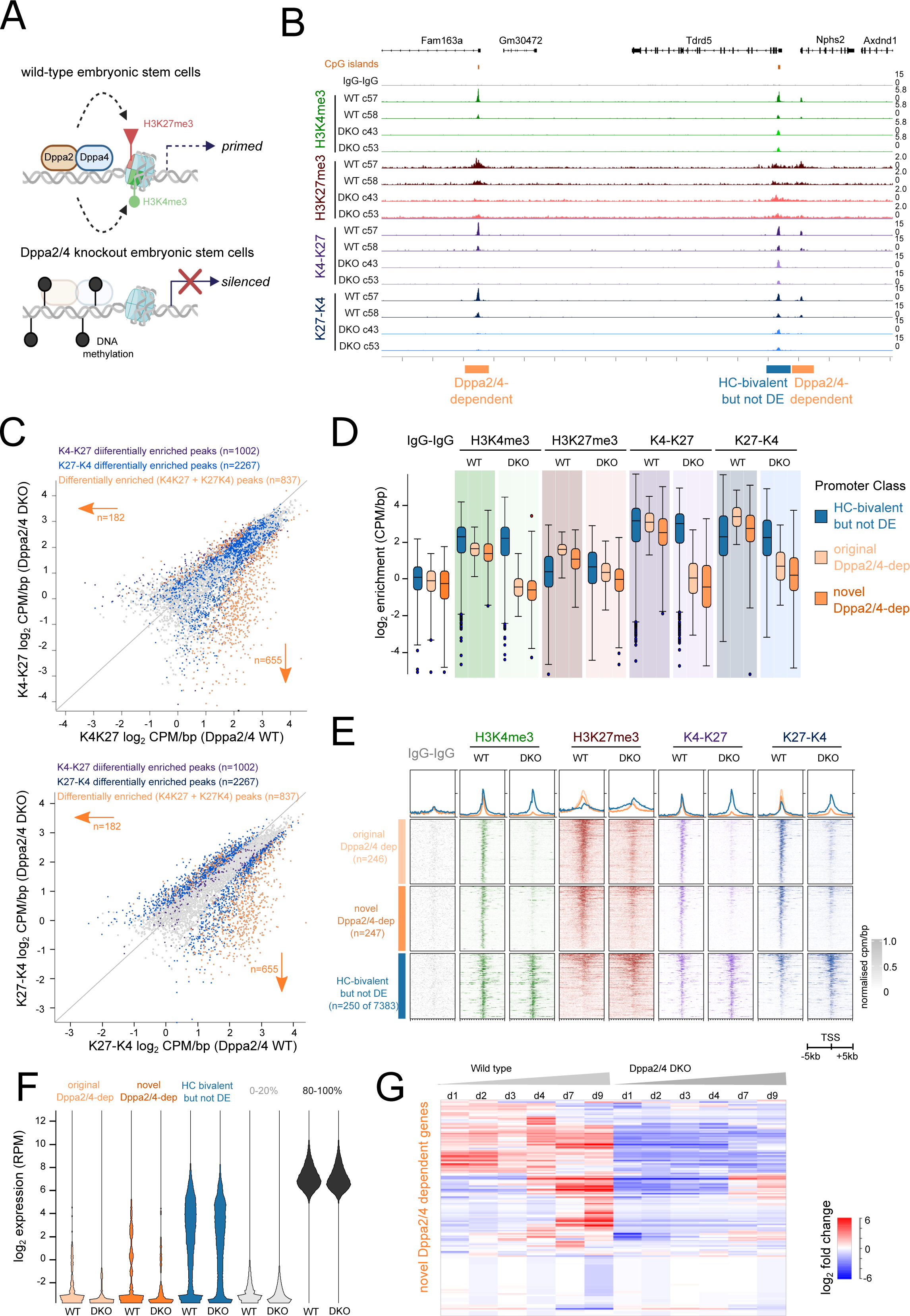
Profiling bivalent chromatin dynamics in DPPA2/4 knockout mouse embryonic stem cells. (A) Schematic depicting how Dppa2/4 maintain both H3K4me3 and H3K27me3 at a subset of bivalent genes, priming them for future activation. Loss of Dppa2/4 leads results in loss of both H3K4me3 and H3K27me3 and gain of repressive DNA methylation. (B) Genome browser view of wild type (WT, dark) and Dppa2/4 double knockout (DKO, light) embryonic stem cell clones. Two clones of each genotype are shown. In-line total H3K4me3 (green), total H3K27me3 (red) and bivalent K4-K27 (purple) and K27-K4 (blue) reChIP data tracks are shown. Dppa2/4 dependent promoters (loose bivalency when Dppa2/4 absent) are denoted by orange bars. (C) Scatterplots showing enrichment (log_2_ CPM/bp) for K4-K27 (top) and K27-K4 (bottom) reChIPs between wild type (x-axis) and Dppa2/4 double knockout (DKO) (y-axis) across all gene promoters. Highlighted are those differentially enriched in the K4-K27 (purple), K27-K4 (blue) or both (orange) reChIP datasets. (D) box plot showing normalised enrichment (CPM/bp) of previously annotated Dppa2/4-dependent genes (light orange) and novel Dppa2/4-dependent genes (dark orange) across the different datasets and clones. As a comparison a subset of Dppa2/4-independent genes (high-confidence bivalent promoters that do not change) are shown (blue). (E) Enrichment heatmaps showing normalised enrichment of previously annotated Dppa2/4-dependent genes (top, light orange) and novel Dppa2/4-dependent genes (middle, dark orange) across the different datasets averaging across clones. As a comparison a subset of Dppa2/4-independent genes (bivalent promoters that do not change) are shown (bottom, blue). (F) Log_2_ RPM expression levels of original (light orange), novel (dark orange) Dppa2/4 dependent genes and high confidence but not differentially enriched (blue) genes across the different datasets between wild type (WT) and Dppa2/4 double knockout (DKO) cells. As a comparison the bottom 20% (light grey) and top 20% (dark grey) expressed genes are shown. (G) log_2_ normalised expression levels of novel Dppa2/4 dependent genes during 9 days of mouse embryoid body differentiation in wild type cells (left) and Dppa2/4 double knockout cells (right). Each gene has been normalised separately across the time series to aid visualisation of expression patterns.

Given the improved sensitivity of our method we next sought to determine whether there may be more widespread changes in chromatin bivalency in Dppa2/4 knockout ESCs compared to what had been previously reported (3,4). Firstly, we called peaks for the reChIP samples. This revealed 13,813 peaks that were bivalently marked in either wild-type and/or Dppa2/4 DKO cells in both reChIP directions. We then classified the bivalent peaks as above using the in-line total H3K4me3 and total H3K27me3 ChIP datasets. This gave 6,146 high-confidence bivalent peaks (i.e. enriched in both H3K4me3, H3K27me3 and both reChIP orientations) in either WT and/or Dppa2/4 DKO cells. To determine if any peaks were gained or lost specifically in Dppa2/4 DKO cells, we performed differential enrichment test using EdgeR (see methods). There were 2,267 and 1,002 differentially enriched bivalent regions in the K27-K4 and K4-K27 reChIP datasets respectively, of which 837 were shared (Figure 4C). This included promoters previously described as Dppa2/4-dependent (4), but also novel bivalent regions detected using the increased sensitivity of our method. Consistent with previous results (3,4), the majority of these were downregulated or absent in the Dppa2/4 DKO cells (Figure 4C).

Promoter-centric analysis revealed differential enrichment of both bivalent K4-K27 and K27-K4 at 493 gene promoters (Figure 4D) which is approximately 2-fold more than what had previously been reported (4). The newly identified (novel) Dppa2/4 dependent promoters had similar levels of enrichment of total H3K4me3, total H3K27me3 and K4-K27 and K27-K4 reChIPs compared to previously known (original) Dppa2/4 dependent promoters (Figure 4D, E). Both original and novel Dppa2/4-dependent bivalent promoters were similarly expressed in undifferentiated ESCs (Figure 4F). Moreover, similar to original Dppa2/4 dependent promoters (4), the novel Dppa2/4 dependent promoters also failed to be upregulated upon embryonic stem cell differentiation (Figure 4G). In summary, this proof-of-principle experiment supports the ability of our method to detect dynamic changes in bivalent chromatin landscapes with high sensitivity and resolution.

## Discussion

Here we present a refined low-input sequential ChIP-reChIP method to robustly and accurately map bivalent chromatin genome-wide. Compared to previously published methods and datasets our approach has several advantages. Firstly, the method requires a substantially reduced input number of cells with just 2 million cells sufficient to generate high quality H3K4me3-H3K27me3 and H3K27me3-H3K4me3 reChIP datasets along with in-line total H3K4me3, total H3K27me3 and IgG-IgG controls. This is a dramatic improvement from the typically 10 million cells or more needed per dataset in other methods (10,14,19) and will facilitate the investigation of these domains in samples where cell numbers are limiting. Next, the data generated has very clear peak enrichments with low background signal enabling standard peak-calling and bioinformatic pipelines to be used to call and classify bivalent regions. While we optimised this method in mouse embryonic stem cells, we envisage its widespread applicability in many different cell lines and tissues.

A key step in all chromatin immunoprecipitation experiments is generating high quality mononucleosomes. The method presented here uses MNase digestion, however we have successfully performed bivalent reChIP experiments from similar number of cells using sonication to shear chromatin with very similar results (data not shown). Importantly MNase/sonication conditions must be optimised for each cell type to ensure predominantly mononucleosome distribution. Over-digested chromatin may not perform well in immunoprecipitation reactions, while under-digested chromatin will confound downstream analysis as it decreases the genomic resolution that can be analysed. In our protocol we implemented a pre-clearing step and found that this drastically improved the signal-to-noise in our experiments. By pre-incubating chromatin with dynabeads, non-specific binding of chromatin fragments to the beads is reduced, removing background and facilitating lower input amounts. Our protocol uses many wash rounds following the immunoprecipitation reactions. We found these to be critical to achieve low background levels. Lastly, we also tested different elution conditions and found that while both SDS-based and peptide-elution approaches behaved similarly, peptide competition elution had higher background levels at long incubation times. Either method could be used in reChIP protocols, however due to cost and availability we opted for SDS-based elution followed by buffer exchange and chromatin concentration prior to the second immunoprecipitation reaction.

Controls are an important part of any experimental design. The IgG-IgG reChIP control provides an estimation of the level of background non-specific binding. When possible, we recommend running a diagnostic qPCR for known bivalent regions and controls prior to library preparation and sequencing. If the IgG-IgG reChIP pulldown amounts are high by qPCR analysis this often indicates the reChIP experiment has not performed well. The IgG-IgG can also be used as normalisation for peak calling in addition to or instead of input samples. Perhaps the most critical control is to perform the reChIP experiment in both orientations (H3K4me3 followed by H3K27me3 and *vice versa*). Our analysis revealed several thousand peaks that are detected in one but not the other reChIP dataset, likely due to the first immunoprecipitation signal carrying through non-specifically from the first immunoprecipitation into the second. This is a common caveat in sequential ChIP experiments and extremely hard to completely eliminate. Therefore, to control for this, any reChIP experiment should always be performed in both orientations to be sure that the detected peaks are indeed due to the presence of both marks on the chromatin.

We also explored other controls that have been used by other studies. One commonly used control is to perform the first immunoprecipitation using H3K4me3 or H3K27me3 and then follow this with a second immunoprecipitation using IgG (19). The rationale behind this is that IgG is non-specific and so there should be no final overall enrichment. However, in our experience, we found that H3K4me3-IgG or H3K27me3-IgG reChIPs mirrored the first immunoprecipitation (data not shown). IgG immunoprecipitation will randomly sample from the pool of chromatin and so if performed as the second immunoprecipitation, this will subsample the already enriched H3K4me3 or H3K27me3 pool of chromatin. Consequently, we have not found this to be a useful control in our experiments or analyses.

When assessing bivalent chromatin, many studies have performed *in silico* merges of independently derived H3K4me3 and H3K27me3 datasets. Theoretically this is unable to distinguish between *bone-fide* bivalent chromatin from allelic or sample heterogeneity. Previous studies in human T-cells (10) have suggested that as much as 14% of bivalent regions called using this approach are false-positives and not true bivalency. Similarly, previous studies in mouse ESCs revealed 1,661 (24%) of 6,817 *in silico* merge bivalent regions were not captured by sequential reChIP (14). In our data, almost all (97%) bivalent regions called using the *in silico* merge approach were also classified as bivalent in our reChIP data. Thus our method is able to capture all predicted bivalent regions suggesting that in mouse embryonic stem cells either approach is sufficient to analyse bivalent chromatin. However we also reveal thousands of additional K4-biased, K27-biased and low confidence bivalent regions. Whether this is the case in other cell types remains unknown and it remains highly likely that reChIP is still required to profile other cell types with stable heterogeneity or complex samples containing multiple cell types and states.

In summary we present a detailed highly optimised method to accurately detect bivalent chromatin dynamics from just 2 million cells. Our protocol uses readily available reagents and equipment found in most molecular biology laboratories and can be adapted to profile this unique form of epigenetic plasticity in any cellular context with the confidence that any conclusions are free from potential confounding effects of cellular heterogeneity.

## Conclusions

Our refined sequential reChIP method provides a useful resource for the wider epigenomics and chromatin biology fields. The optimised protocol accurately and robustly detects twice as many bivalent regions in mouse embryonic stem cells as previously identified (14), from as little as 2 million cells. Consistent with current models, the bivalent regions occur predominately at CpG-rich promoters that are dynamically regulated during differentiation. Lastly our analysis of Dppa2/4 knockout cells confirms the ability of our method to detect changes in the bivalent chromatin landscape. This method will facilitate accurate profiling of the dynamics of bivalent chromatin in other contexts, greatly improving our understanding of this unique form of epigenetic plasticity.

## Methods

### Cell culture

Mouse embryonic stem cells were cultured on feeder-free gelatinised plates at 37°C, 5% CO_2_ using standard serum/LIF conditions (high-glucose DMEM supplemented with 15% fetal bovine serum, 1x GlutaMax, 1x penicillin, 1x streptomycin, 0.1mM nonessential amino acids, 50mM beta-mercaptoethanol and LIF (made in house in HEK293 cells and titrated for optimal ESC growth)). Cells were regularly tested for mycoplasma contamination using the Mycoplasma PCR Detection Kit (abcam ab289834). E14 mouse embryonic stem cells were a gift from W. Reik’s laboratory. Wild type and Dppa2/4 double knockout clones were generated in (4,23) and cultured as above. Cells were not authenticated. Cells were cultured at least 2 passages from thawing prior to chromatin collection. Biological replicates were collected from different passages on separate days.

### Cell collection and fixation

Cells were seeded on multiple plates and grown to near-confluency. At time of harvest one plate was used to determine cell concentration. Cells on remaining plates were washed with PBS and fixed with 1% methanol-free formaldehyde (Thermo Scientific 28908) in DMEM at room temperature for 8 minutes, quenched with 0.125M glycine and scraped off cell culture dishes. Cell slurry was washed with ice-cold PBS, resuspended in PBS/EDTA, aliquoted to 2×10^6^ cells per vial, spun down and snap frozen on dry ice for storage at −80°C. Cell pellets were used within 6 months of collection.

### Sequential chromatin immunoprecipitation

Pellets of 2×10^6^ cells were lysed with 100μl NP buffer (10mM TrisHCl pH7.4, 1M sorbitol, 50mM NaCl, 5mM MgCl_2_, 0.075% IGEPAL) freshly supplemented with 0.385mM beta-mercaptoethanol (Gibco 21985-023) and 1.8mM spermidine (Sigma 05292) on ice. Chromatin was digested using 2.4μl per sample of MNase (NEB) for 37°C for 15 minutes with gentle shaking at 600rpm. Reactions were stopped with 26.4μl STOP buffer (50mM EDTA, 0.5% TritonX-100, 0.5% sodium deoxycholate), incubated on ice for >5 minutes, vortexed and sample diluted to 580μl in ChIP buffer (20mM TrisHCl pH8.0, 2mM EDTA, 150mM NaCl, 0.5% Triton X-100) containing protease inhibitor cocktail (cOmplete EDTA-free Protease Inhibitor Cocktail, Roche). Chromatin was precleared by adding 20μl prewashed Protein A dynabeads (Invitrogen 10002D) and incubating at 4°C on rotator for >2 hours. 5% of the sample was set aside as input, the remaining chromatin was divided amongst separate tubes containing protein A dynabeads pre-incubated with either 2μl anti-H3K4me3 (Millipore 07-473), 10μl anti-H3K27me3 (CST 9733) or 1μg IgG (Invitrogen) antibodies. First immunoprecipitation was performed overnight at 4°C with rotation. Antibody-chromatin complexes were washed 3x in low salt buffer (20mM TrisHCl pH8.0, 2mM EDTA, 150mM NaCl, 1% Triton X-100, 0.1% SDS), 3x in high-salt buffer (20mM TrisHCl pH8.0, 2mM EDTA, 500mM NaCl, 1% Triton X-100, 0.1% SDS), 2x in LiCl buffer (0.35M LiCl, 1% IGEPAL, 1% sodium deoxycholate, 1mM EDTA, 10mM Tris-HCl pH7.5) and 2x in TE on ice. Complexes were eluted in 100μl elution buffer (10mM TrisHCl pH8.0, 1mM EDTA, 1% SDS) containing fresh protease inhibitor cocktails for 30 minutes at 37°C with shaking. 10% sample was set aside as total in-line control ChIPs. To dilute SDS volume was increased to 300μl with ChIP buffer containing protease inhibitor cocktails and purified using Amicon Ultra-0.5ml 3KDa filter columns (Millipore) according to manufacturers instructions, recovering approximately 50μl chromatin per IP reaction. The second immunoprecipitation was performed using the alternate antibody or IgG control overnight at 4°C with rotation and chromatin washed and eluted as previously. In-line control, input and ChIP samples were heated at 65°C for 2.5 hours to reverse cross-links, treated with RNAseA (NEB) for 30 minutes at 37°C, proteinase K (NEB) for 1 hour at 37°C, and purified using Ampure beads (Beckman Coulter) at a 1:1.8 ratio.

### Peptide elution experiments

Peptide elution experiments were performed by resuspending washed dynabead-antibody-chromatin complexes in 200μl peptide elution buffer (50mM Tris-HCl pH8.0, 5mM EDTA, 100mM NaCl, 0.5% sodium deoxycholate, 0.1% SDS) supplemented with protease inhibitors containing 10μg/ml H3K4me3 (abcam ab1342) or H3K27me3 peptides (abcam ab1782) on rotator at 4°C for 3 hours or overnight. For the IgG control sample 10μg/ml of a 1:1 mix of H3K4me3 and H3K27me3 peptides was used.

### qPCR analysis

qPCR analysis of purified ChIP DNA was performed in technical duplicate for each primer pair using 2x SYBR mastermix (Applied Biosystems Cat#4385612) according to manufacturer’s instructions in a 6-10μl reaction using the primer sequences as below.

H3K4me3-only controls

Dppa2_forward GCCAAACACAGACTACGCTA
Dppa2_reverse AACCTACACTATTTTCGCCAGGAT
Dppa4_forward TTCTCAAGATGGAGACTGCTGG
Dppa4_reverse TGGCTATACTCAAAAATGAGGGGC

H3K27me3 only controls

Gm6116_forward GCGGTGAGTACTCTGCTCAA
Gm6116_reverse CCATCCAGTACTGTGGGCTC
K27me_R1_forward TGCCTGCAATTCGTCCTCTT
K27me_R1_reverse ACGAAGCAGCCGTGTAAGAA

Bivalent regions

Csf1_forward GAGCACCGAGGCAAACTTTC
Csf1_reverse GAGCCAGGGTGATTTCCCAT
Lmo1_forward AAGCGGGCTCTAATTACCCG
Lmo1_reverse CTGCGAAGTGCTTCACTCCT
Pou4f1_forward CAAAGTGAGGCTGCTTGCTG
Pou4f1_reverse GCGGACTTTGCGAGTGTTTT
Sox6_forward CGATACAGAAGCGCAGGCTA
Sox6_reverse AGGGGCCCTTGTAGATGGAT

### library preparation

Sample DNA concentrations were quantified using Qubit and libraries prepared using NEBNext Ultra II DNA library preparation kit (NEB) according to the manufacturer’s instructions with the following modifications. To achieve optimal final DNA concentrations, samples were re-quantified on the Qubit 3.0 following PCR amplification and an additional 3 cycles (if concentration close to 20ng/μl) or 5 cycles (if concentration << 20ng/μl) performed if needed to obtain the ideal final library concentration of 20-100ng/μl. A maximum of 20 cycles was used for any sample. Note that due to the low amounts of DNA obtained from the protocol, concentration measurements prior to amplification typically occur at the lower limit of detection and even if zero values are obtained, libraries can often still be generated. Libraries were purified using NEBNext sample purification beads and checked using Agilent Tapestation 2200 or 4150 on a high-sensitivity tape (HSD1000) aiming for a final library with dominant peak size of 270bp. Libraries were pooled and sequenced using the Illumina NextSeq500 platform with a target read depth of 20 million SE75bp reads per sample.

### Data pre-processing

Single-end reads in fastqs were trimmed and filtered for quality (phred33 score > 20) and length (>20bp) using TrimGalore (https://github.com/FelixKrueger/TrimGalore) v0.6.6 in single-end mode. Trimmed and quality filtered reads were then aligned to the mm10 mouse genome (GRCm38.p6) using bwa-mem(24) (bwa v0.7.13) with default parameters. Alignments were then converted to the bam format and indexed using samtools v1.9. Duplicate alignments were then marked with using *MarkDuplicates* (picard v2.6.0, https://broadinstitute.github.io/picard/) and re-indexed with samtools(25). bigWig files containing CPM/bp normalised coverage values for each sample were derived from duplicate marked bam files using *bamCoverage* (DeepTools v3.5.0 (26)) whilst excluding ENCODE blacklisted genomic regions (27) for the mm10 genome (v2).

### ChIP and reChIP data analysis

Aligned read (bam) files were imported into SeqMonk software version (v1.48.1) (http://www.bioinformatics.babraham.ac.uk/projects/seqmonk) for all downstream analysis using standard parameters (no deduplication, MAPQ>20, primary alignments only), extending reads by 200 bp. In-line total H3K4me3, in-line total H3K27me3, IgG-IgG, bivalent K4K27 and bivalent K27K4 datasets for E14 ESCs, Dppa2/4 WT clones and Dppa2/4 DKO clones were generated in this study. Single-ChIP-seq data from 10 million cells of H3K4me3, H3K27me3 and input controls were obtained from (4) (GSE135841).

Peak calling for in-line total H3K4me3, total H3K27me3 and K4-K27 and K27-K4 reChIP datasets was performed using the two biological replicates and IgG-IgG as input control using the inbuilt MACS peak caller (p-value<1e-5, fragment size 300). For total H3K4me3 and total H3K27me3 data derived from 10 million cells, input DNA was used in peak calling. Peaks from different datasets were merged together and overlapping peaks or those within 200bp were stitched together. Differential enrichment was performed using edgeR (p-value cut-off 0.05 with Benjamini-Hochberg corrections for multiple testing applied). Normalised read densities within peaks were calculated as log_2_-transformed read counts in peaks corrected for library size (counts per million reads) and peak length (per bp) yielding CPM/bp. Promoters were defined as the region spanning 1kb upstream and 1kb downstream of the transcription start site of genes. For each peak set, the fraction of reads in peaks (FRiP) were calculated by first counting the reads in each bam at each peak with the csaw R package (v1.28.0) and then dividing the sum of these counts by the total number of reads in the library. Alluvial plots demonstrating the differences in bivalent peak annotations were plotted using the ggalluvial R package (v0.12.5).

### Gene expression analysis

Gene expression data was obtained from (4) (GSE135841). Data was trimmed with Trim Galore (v0.4.4, default parameters) and mapped using HiSat2 v2.1.0 to the mouse GRCm38 genome assembly. RNA-sequencing analysis was performed using SeqMonk software using inbuilt RNA-sequencing quantification pipeline. Expression values represent log_2_ transformed quantification of merged transcripts counting opposing-strand reads over exons. Gene expression heatmaps are normalised for each transcript independently by subtracting the median value for that transcript across all samples from each sample value. Bean plots represent smoothed density of all points over the bandwidth window corresponding to 5% of the total quantitation range displayed in the plot.

### Genomic enrichment heatmaps and trackplots

To generate genomic enrichment heatmaps and trackplots bigWigs containing CPM/bp normalised read densities were used. For heatmaps, each peak region was first extended by 5kb upstream and downstream and then split into 100 equally sized bins using the GenomicRanges R package (v1.46.1) (28). The average CPM/bp was calculated for each bin for each ChIP using the EnrichedHeatmap R package (v1.24.0) (29). Bins with values surpassing the 99th percentile of all bins within each ChIP were masked (i.e. assigned the 99th percentile value) to eliminate extreme outliers from affecting colour scales. Each bin was then scaled relative to the highest value (so values range between 0-1 and represent the relative enrichment of signal across all regions). Enriched heatmaps were then plotted using the same package, with the average bin value plotted as continuous curves atop each heatmap. Genomic track plots were plotted using the rtracklayer (v1.54.0) (30) and Gviz R packages (v1.38.4) (31). CpG island annotations for the mm10 genome were retrieved from the UCSC genome browser (32).

### Gene ontology

The enrichment of gene ontologies across subclasses of genes with bivalent promoters (HC, K4,K27,LC) were determined using the clusterProfiler R package (v4.2.2) (33). Gene symbols were first converted to entrez id’s using the biomaRt R package (v2.50.3) (34) and were input alongside a background list of all expressed genes to clusterProfiler using the *compareCluster* function against the Gene Ontology (GO) database (35). Significantly enriched GO terms were those with a Benjamini-Hochberg (BH) corrected p-value < 0.05, had atleast 10 genes present in the pathway and a gene ratio (genes in subclass/genes in pathway) > 0.01. Representative pathways were plotted using the ggplot2 R package (v3.3.5) (36).

### Motif analysis

Enrichments for transcription actor binding motifs in peak subclasses were calculated using the monaLisa R package (v1.0.0). Position weight matrices for transcription factor binding sites in vertebrates were retrieved from the JASPAR2020 database (37). Binned motif enrichment for peak subclasses (HC, K4, K27, LC) were then conducted in monaLisa (38) while including randomised sequences modelled off all bivalent peaks (made with the regioneR R package (v1.26.1) (39)) as the background. Significant enrichments were those with a BH-adjusted p-value < 0.05 and log2-fold enrichment over random sequences > 1. Motif heatmaps were also plotted using monaLisa.

### CG-content

CG content was determined for peak subclasses (HC, K4, K27, LC) as well as the same random control sequences above using the Biostrings R package (v2.62.0) (40) by first calculating all oligonucleotide frequencies and then by summing C and G frequencies. All dinucleotide frequencies were calculated using monaLisa with the *plotBinDiagnostics* function and then GC/CG dinucleotide frequencies were summed. These data were plotted using ggplot2, and the significance of comparisons were determined using pairwise t-tests followed by BH-adjustments of p-values to account for multiple comparisons.

### Chromatin state discovery

bam files were first converted to the bed format using bedtools (*bamtobed*) (v2.27.1) (41). bed files were then partitioned into 200bp bins and then binarized for the determination of bin-specific enrichments (providing replicate in-line ChIPs or reChIPs and using IgGIgG as the control) using ChromHMM (*BinarizeBed*) (v1.24) (42). Hidden Markov Models were then used to discover chromatin states across these genomic bins using ChromHMM (*LearnModel*) using a 5-state model. Segment bed files containing chromatin state annotations were then overlapped with our bivalent peak annotations (HC, K4, K27, LC), where each peak was then re-assigned to the chromatin state with the highest degree of overlap using the GenomicRanges R package (*findOverlaps* and *pintersect*) (v1.46.1) (28). Heatmaps containing emission probabilities, transition probabilities, TSS enrichments and annotation overlaps from the ChromHMM model were then plotted using the ComplexHeatmap R package (v2.10.0) (43).

### Software

Plots were generated using SeqMonk software (v1.48.1) or R (v4.1.2/RStudio v2022.02.0+443), and edited in Inkscape. Schematic figures were made with BioRender.com with publishing licence agreement numbers *RM25UDOLG3, UQ25UDOE0U* and *MC25UDOPHO*.

## Supplemental Figures and Tables

**Supplemental Figure 1.**
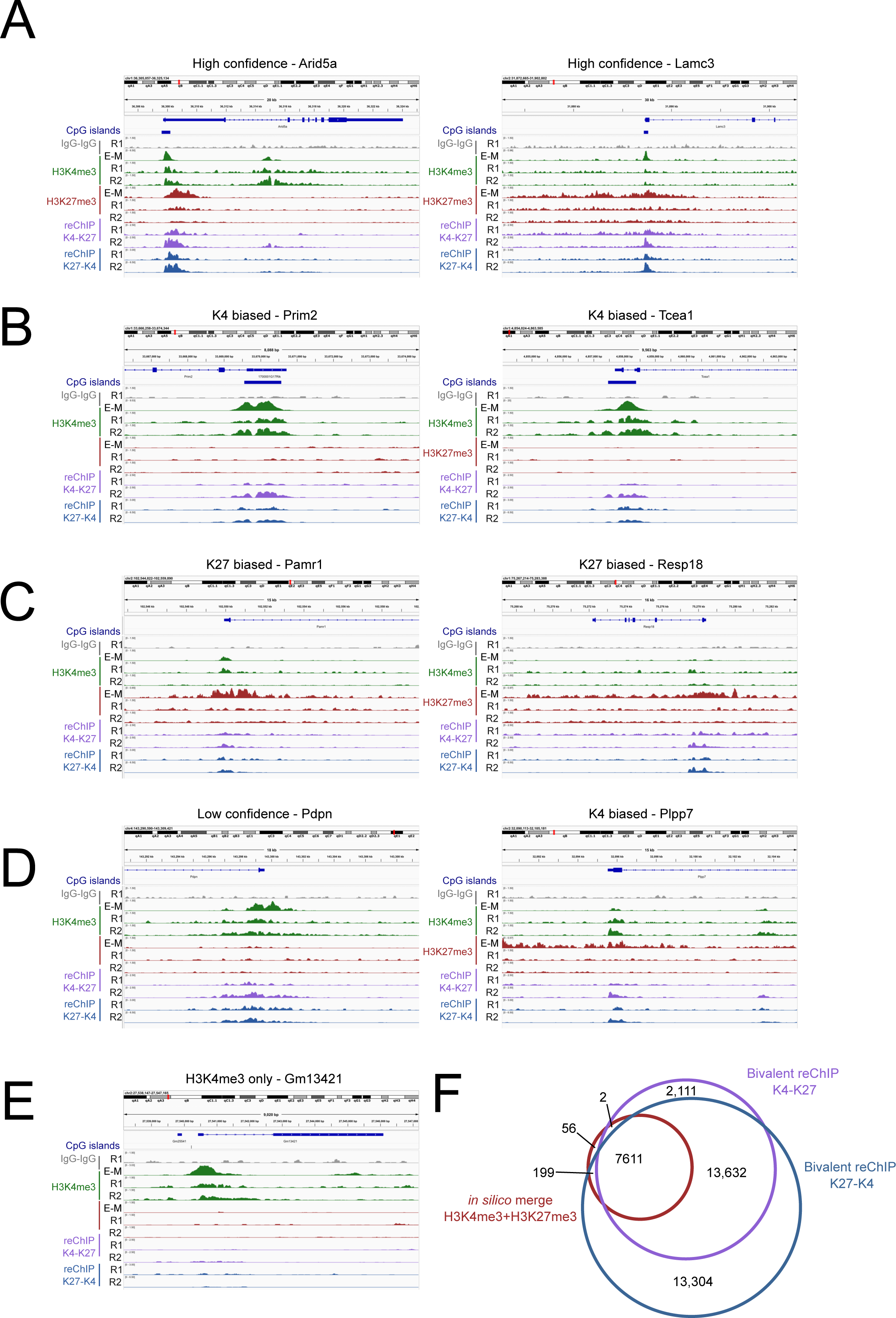
, related to Figure 1 and 2. (A-E) Genome browser views of high-confidence (A), K4-biased (B), K27-biased (C), low confidence (D) and H3K4me3-only (E) genes showing H3K4me3 (green), total H3K27me3 (red) and K4-K27 (purple) and K27-K4 (blue) reChIP datasets. Height of peak represents CPM/bp. E-M represents data from independent 10 million cell total H3K4me3 and total H3K27me3 (GSE135841) (4). R1 and R2 are two independent biological replicates from this study. (F) Venn overlap between peaks classified using in silico merge of independent 10 million cell total H3K4me3 and total H3K27me3 (GSE135841) (4) (red) versus peaks called with K4-K27 (purple) and K27-K4 (blue) reChIPs (this study). Note numbers are slightly different to those in Figure 3A as the total number of peaks was summed across bivalent reChIP and total ChIP datasets (as opposed to just bivalent reChIP in Figure 3A).

## Declarations

### Ethics approval and consent to participate

Not applicable

### Consent for publication

Not applicable

### Availability of data and materials

The datasets generated during the current study have been deposited in the short read archives (SRA) and gene expression omnibus (GEO) under the accession GSE242686. Gene expression data and previous ChIP-seq data was obtained from (4) (GSE135841).

### Authors’ contributions

M.A.E.-M. conceived, designed and supervised the study, performed experiments, analysed data and wrote the paper. W.H. optimised and performed reChIP experiments and generated libraries. J.S. processed data and performed data analysis. E.G. independently verified the reChIP method and wrote the detailed protocol.

## Supporting information

Supplemental_File1_DetailedProtocol

## Acknowledgements

We would like to thank all past and present members of the Eckersley-Maslin laboratory for their feedback throughout the project. Mouse E14 embryonic stem cells were kindly provided by Wolf Reik’s laboratory. We thank Billy Hamilton for assistance with synthesising LIF in house. We thank Gisela Mir Anau, Tim Semple and Stuart Craig from the PeterMac Molecular Genomics Core facility for advice in library preparation of low input samples and high-throughput sequencing runs. Our laboratory is located on the lands of the Wurundjeri people of the Kulin Nation and we pay our respects to their elders, past present and emerging, and recognise their continuing connection to country and community.

