## Supplemental_File1_DetailedProtocol for "A low-input high resolution sequential chromatin immunoprecipitation method captures genome-wide dynamics of bivalent chromatin"

**This is a detailed protocol for performing sequential ChIP-reChIP to map H3K4me3-H3K27me3 bivalent chromatin regions. It has been optimised using mouse embryonic stem cells and so may need to be refined based on your cell type of interest. For more details please refer to the accompanying manuscript (Ho *et al*. 2023).**

### General workflow

A. Chromatin fixation (2 hours)
B. Chromatin fragmentation and Primary antibody incubation (2 hours hands on time with overnight incubation)
C. First elution, buffer exchange and secondary antibody incubation (2 hours hands on time with overnight incubation)
D. Secondary elution, de-crosslinking, and DNA purification (4 hours)

**Note:** As a standard throughout the protocol, we recommend using low bind tubes and tips, and advise that all buffers be prepared using RNase and DNase free reagents. Furthermore, while most commercially available antibodies will have a recommended volume to use, we recommend validating and titrating antibody volumes and cell numbers prior to starting. Finally, we recommend designing and testing quantitative PCR primers to amplify positive and negative control regions, ensuring they have good efficiencies and melt-curves. PCR products should be between 80-120 base pairs. Validated primer sequences for mouse Embryonic Stem Cells can be found in the accompanying manuscript.

### Key resource & reagent table

| Reagent or Resource | Source | Catalogue No. |
|---|---|---|
| 16% Formaldehyde (w/v), Methanol free | Life Technologies | 28908 |
| Glycine | Sigma | G8898 |
| Sodium deoxycholate | Sigma | 30970 |
| Lithium Chloride solution 8M | Sigma | L7026 |

| | | |
|---|---|---|
| MNase | NEB | M0247S |
| RNaseA | NEB | T3018-2 |
| ProteinaseK | NEB | P8107S |
| Protein A DynaBeads | Thermo Fisher | 10002D |
| Protein LoBind 1.5ml tubes | Eppendorf | 0030108442 |
| Magnetic Rack | Invitrogen | 12321D |
| Protease Inhibitor cocktail | Roche | 05892791001 |
| Amicon Ultra | Millipore | MPUFC5003BK |
| NEBNext Ultra II DNA Library Prep Kit | NEB | E7645L |

**<u>Prepare buffers as outlined below</u>**

- NP buffer
- Complete Chromatin Immunoprecipitation buffer
- Low salt wash buffer
- High salt wash buffer
- LiCl wash buffer
- Elution buffer

**NP buffer**

| Reagent | Final Concentration | Amount in 100ml |
|---|---|---|
| 1M Tris pH7.4 | 10mM | 1ml |
| Sorbitol | 1M | 18.217g |
| 1M NaCl | 50mM | 5ml |
| 1M MgCl2 | 5mM | 0.5ml |
| 1M CaCl2 | 1mM | 0.1ml |
| IGEPAL | 0.075% | 75µl |

| | | |
|---|---|---|
| ddH2O | | Up to 100 ml |

**Chromatin Immunoprecipitation Buffer**

| Reagent | Final Concentration | Amount in 500ml |
|---|---|---|
| 1M Tris pH7.4 | 20mM | 10ml |
| 0.5M EDTA | 2mM | 2ml |
| 1M NaCl | 150mM | 75ml |
| Triton X100 | 0.1% | 0.5ml |
| ddH2O | | Up to 500ml |

**Low salt wash buffer**

| Reagent | Final Concentration | Amount in 500ml |
|---|---|---|
| 1M Tris pH8 | 20mM | 10ml |
| 0.5M EDTA | 2mM | 2ml |
| 5M NaCl | 150mM | 15ml |
| Triton X100 | 1% | 5ml |
| 10% SDS | 0.1% | 5ml |
| ddH2O | | Up to 500ml |

**High Salt wash buffer**

| Reagent | Final Concentration | Amount in 500ml |
|---|---|---|
| 1M Tris pH8 | 20mM | 10ml |

| | | |
|---|---|---|
| 0.5M EDTA | 2mM | 2ml |
| 5M NaCl | 500mM | 50ml |
| Triton X100 | 1% | 5ml |
| 10% SDS | 0.1% | 5ml |
| ddH2O | | Up to 500ml |

**LiCl wash buffer**

| Reagent | Final Concentration | Amount in 500ml |
|---|---|---|
| 8M LiCl | 250mM | 15.625ml |
| IGEPAL | 1% | 5ml |
| 10% deoxycholate | 1% | 50ml |
| 0.5M EDTA | 1mM | 1ml |
| 1M Tris pH7.4 | 10mM | 5ml |
| ddH2O | | Up to 500ml |

**Elution Buffer**

| Reagent | Final Concentration | Amount in 10ml |
|---|---|---|
| 1M Tris pH7.4 | 10mM | 0.1ml |
| 0.5M EDTA | 1mM | 0.02ml |
| 10% SDS | 1% | 1ml |
| ddH2O | | Up to 10ml |

### Section A: Chromatin fixation

*Before you start: pre-warm DMEM; pre-cool centrifuge to 4 degrees; add Roche cOmplete EDTA-free protease inhibitor tablets to PBS/EDTA and keep on ice until required.*

*Note: all steps must be performed on ice or at 4 degrees unless otherwise specified.*
*Note: chromatin fixation can also be performed on cells in suspension by resuspended a pellet of a known number of washed live cells directly into the 1% formaldehyde solution and incubating for 8 minutes on a rocker, quenching with glycine as below and then mild centrifugation to get a fixed cell pellet( rather than scraping cells).*

1. Grow cells until they are 70-80% confluent across several plates. One will be used for counting and the remaining for fixation (collection plates).
2. Collect and count cells on the counting plate: Remove media from the counting plate and wash once with DPBS.
3. Add 1ml of Trypsin or appropriate dissociation reagent per 10cm plate and incubate until cells lift off plate as single cells.
4. Quench trypsin with 5ml of media.
5. Take aliquot and count cells to determine total cell number per plate and therefore the total number of cells overall.
6. Prepare 1% formaldehyde solution by adding 0.625ml of 16% formaldehyde solution to 9.375ml of prewarmed DMEM per 10cm collection plate. Scale up volumes if you have more than one 10cm collection plate.
7. Remove media and wash cells on collection plates with DPBS.
8. Cross link cells by adding 10ml of 1% formaldehyde solution per plate of adherent cells for 8 min at room temperature.
9. Quench with 1ml 1M glycine per plate of adherent cells to a final concentration 125mM, for 5 min at room temperature.
10. Pour off medium and wash cells with 10ml of ice cold DPBS.
11. Pour off DPBS, scrape cells in residual PBS and transfer to low bind 1.5ml tube. Pool cells from all plates into one tube.
12. Centrifuge for 10min at 500g at 4 degrees.

13. Remove supernatant and resuspend cells to 2x10^7 cells per 1ml in PBS/5mM EDTA containing protease inhibitors. Aliquot by adding 0.1ml cell slurry (corresponding to 2x10^6 cells) per low-bind tube.
14. Pellet aliquots by centrifugation for 5 minutes at 500g and 4 degrees.
15. Remove supernatant and snap freeze on dry ice or in liquid nitrogen. Store pellets at -80 degrees for up to 6 months.

### Section B: Chromatin fragmentation and primary antibody incubation - Day 1

*Before you start: pre-cool centrifuge; add Roche cOmplete EDTA-free protease inhibitor tablets to ChIP buffer and NP buffer and keep on ice until needed.*

*Note: Unless otherwise specified all procedures should be carried out between 2-8 degrees*
*Note: Add protease inhibitor cocktail to sufficient volumes of NP buffer and ChIP buffer.*
*Note: NP buffer composition and cell lysis conditions may need to be optimised depending on your cell type.*
*Note: volumes below are to process one sample corresponding to one vial of 2x10^7 cells and will result in input, in-line H3K4me3, in-line H3K27me3, IgG-IgG reChIP and bivalent K4-K27 and K27-K4 reChIP samples that can be further processed by qPCR and/or library preparation for sequencing. If you have more than one sample (e.g. biological replicate or other condition), scale volumes accordingly. We typically do not process more than 4 samples at any time.*
*Note: sonication can be used instead of MNase digestion. We routinely use both fragmentation methods in performing reChIP experiments with similar results.*

**Bead preparation – 30 minutes**

16. Prepare a sufficient amount of Protein A dynabeads for chromatin preclear and antibody binding. For each sample in the reChIP experiment you will need 150µl. This corresponds to:
    - 20µl of beads to preclear 2x10^6 cells.
    - 120µl of beads for antibody-complex formation (20µl per antibody-complex of which there are 6 reactions in total: 2xIgG, 2x H3K27me3, 2xH3K4me3)

- 10μl for pipetting errors

17. Take 150μl Protein A dynabeads, place on magnetic rack and remove supernatant.
18. Remove tube from rack and resuspend in 500μl of cold ChIP buffer containing freshly added protease inhibitor cocktail.
19. Repeat steps 2-3 for a total of 3 washes.
20. After 3 washes resuspend in 150μl of cold ChIP buffer.
21. Keep chilled on ice until needed later.

**Binding Antibody to beads – 15 minutes preparation time plus at least 3 hours incubation**

22. Label 6 low-bind tubes (2X IgG, 2X H3K4me3 and 2X H3K27me3)
23. Add 500μl of cold ChIP buffer to each tube.
24. Add 20μl of pre-washed Protein A dynabeads to each tube.
25. Add appropriate antibody to each tube - 1μg IgG (Invitrogen), 10μl H3K27me3 (CST 9733), 2μl H3K4me3 (Millipore 07-473). If using other antibodies the volumes will need to be titrated to maximise signal:noise. Adding too little antibody will not capture all chromatin containing the modification of interest. Adding too much antibody increases the background non-specific binding.
26. Incubate bead/antibody mix at 4 degrees on a rotator for at least 3 hours.

**Chromatin preparation – 1 hour preparation time plus at least 3 hours incubation**

27. Thaw 1 vial of 2x10^6 crosslinked cells on ice
28. Resuspend cell pellet (2x10^6 cells) in 97.68μl NP buffer supplemented with 0.7μl of 55μM beta-mercaptoethanol, 1.82μl of 0.1M spermidine and freshly added protease inhibitor cocktail.
29. Fragment chromatin with MNase: prepare MNase master mix (20μl per sample)
    - 12μl 10X MNase buffer
    - 1.76μl 100mM DTT
    - 3.84μl dH20
    - 2.4μl MNase
30. Add 20μl of MNase master mix to each tube of 2x10^6 cells in NP buffer.
31. Incubate at 37 degrees with shaking at 600rpm for 7.5 - 15 min.

*Note: This amount of MNase and digestion time will need to be titrated for each cell line. These conditions have been optimised to yield predominantly mono nucleosomal DNA for 2x10^6 mouse embryonic stem cells. If preferred sonication can be used to fragment chromatin instead.*

32. During digestion prepare STOP buffer:
    - 15µl 100mM EDTA
    - 15µl 1%triton/1%deoxycholate solution
33. Add 26.4µl of STOP buffer to each sample to stop the MNase digestion.
34. Incubate on ice for 5 minutes.
35. Vortex each tube for 30 seconds each.
36. Bring volume up to 600µl by adding 473.6µl cold ChIP buffer containing protease inhibitor cocktail.
37. Add 20µl of pre-washed protein A Dynabeads from step 21 to the chromatin and incubate for at least 3 hours at 4 degrees on rotator to pre-clear the chromatin. This is critical to reduce non-specific binding and decrease background signal.

**Overnight incubation with primary antibody – 30 min and overnight**

38. Take all bead-antibody tubes from the 4-degree rotator.
39. Back at the bench place the pre-cleared chromatin sample and 1x IgG, 1 x H3K4me3 and 1 x H3K27me3 bead-antibody complexes per sample on the magnet rack until the solution clears. We recommend sitting the magnetic rack on ice to keep cool during these steps. Alternatively, these steps can be performed in a cold room.
40. The remaining bead-antibody mixes can be stored at 4 degrees until needed on day 2.
41. Take 10µl of pre-cleared chromatin supernatant to a separate tube and label as 5% input control. Keep at 4 degrees until day 3.
42. Remove the supernatant from the antibody-bead complexes on the magnetic rack and discard.
43. Add 200µl of the pre-cleared chromatin supernatant to each antibody/bead mixture.
44. Top up each chromatin/antibody/bead mixture with 300µl of ChIP buffer to final volume of 500µl.

45. Incubate overnight at 4 degrees on a rotator.

### Section C: First elution, buffer exchange and secondary antibody incubation – Day 2

*Before you start: Prepare 30ml of ChIP buffer containing protease inhibitor cocktail. Prepare 5ml of elution buffer containing protease inhibitor cocktail; pre-cool centrifuge to 4 degrees.*

46. Collect chromatin-antibody-bead samples from overnight 4-degree rotation.
47. Wash chromatin-antibody-bead complexes a total of 9 times using the following steps keeping the samples cool by either placing magnet on ice or working in a cold room:
    a. Place tubes on magnets rack and wait for solution to turn clear.
    b. Carefully remove supernatant while on magnet making sure you do not disturb the beads. Make sure you do not let the beads dry out.
    c. Resuspend beads in 500µl of low salt buffer.
    d. Repeat steps a-c for a total of 3x low salt buffer washes, 3x high salt buffer washes, 2x LiCl buffer washes and 2x 1xTE washes.
48. Elute washed complexes in 100µl elution buffer containing fresh protease inhibitor cocktail for 30 min at 37 degrees on a thermomixer. If you do not have access to a shaking heat block, gently flick the tubes periodically during the incubation to ensure beads remain suspended in solution.
49. During elution step prepare 3x Amicon Ultra buffer exchange columns by adding 500µl Milli-Q H20 to columns and spinning at 14000g for 30 min at 4 degrees.
50. After chromatin elution, place samples on magnetic rack and wait for sample to turn clear. Move supernatant containing chromatin to a new low-bind tube.
51. Take 10% volume (10µl) from each chromatin IP as an in-line single ChIP control into a new low-bind tube, label and store at 4 degrees until day 3.
52. Bring each remaining chromatin sample up to 300µl with ChIP buffer containing protease inhibitor cocktail.
53. Decant the H20 from the prepared Amicon Ultra filters and each chromatin sample to a separate filter.
54. Spin at 14000g for 30 min at 4 degrees.

55. Carefully decant flowthrough and discard.
56. Add 500µl ChIP buffer and spin at 14000g for 30 minutes at 4 degrees.
57. Carefully decant flowthrough and discard.
58. Repeat steps 55 and 56 for a total of 2 washes.
59. Recover as much chromatin sample as possible from within the Amicon Ultra filter. (This is usually around 50µl).
60. Bring volume up to 500µl with ChIP buffer containing protease inhibitors.
61. Take antibody bound beads for the second incubation from 4 degrees and place on magnetic rack.
62. Wait for the solution to turn clear and remove the supernatant.
63. Add appropriate chromatin samples to appropriate antibodies:
    - Add the IgG chromatin sample to the IgG-bead complexes.
    - Add the H3K4me3 chromatin to the H3K27me3-bead complexes.
    - Add the H3K27me3 chromatin to the H3K4me3-bead complexes.
64. Incubate overnight at 4 degrees with rotation.

### Section D: Second elution, de-crosslinking and DNA purification – Day 3

65. Wash chromatin-antibody-bead complexes a total of 9 times using the following steps keeping the samples cool by either placing magnet on ice or working in a cold room:
    a. Place tubes on magnets rack and wait for solution to turn clear.
    b. Carefully remove supernatant while on magnet making sure you do not disturb the beads. Make sure you do not let the beads dry out.
    c. Resuspend beads in 500µl of low salt buffer.
    d. Repeat steps a-c for a total of 3x low salt buffer washes, 3x high salt buffer washes, 2x LiCl buffer washes and 2x 1xTE washes.
66. Elute complexes and reverse crosslinks in 100µl elution buffer for a minimum of 2.5 hours up to overnight at 65 degrees on a thermomixer. Note – no protease inhibitors are required for this elution step.
67. Collect the 5% input sample (10µl) and three in-line total control samples (10µl each) from 4 degrees

68. Add 90μl elution buffer to bring the final volume of each control to 100μl.
69. Place at 65 degrees for 2.5 hours on a thermoshaker alongside the reChIP samples to de-crosslink.
70. After de-crosslinking, place all tubes on the magnetic rack. You should have a total of 7 tubes corresponding to input, 3x in-line total controls and 3x reChIPs.
71. When the solution turns clear move supernatant to new low-bind tubes.
72. Add 2μl RNaseA (NEB) to each tube and incubate at 37 degrees for 30 min.
73. Add 2μl Proteinase K (NEB) to each sample and incubate at 37 degrees for 1 hour.
74. Purify DNA using Ampure beads.
    a. Bring Ampure beads to room temperate prior to use.
    b. Add beads to sample in a 1:1.8 ratio (e.g. 180μl beads to 100μl sample) and pipette up and down to mix.
    c. Incubate at room temperate for 5 minutes.
    d. Place tubes on magnetic rack for 5 mins and remove supernatant.
    e. While tubes are on the rack wash the beads with 400μl of freshly prepared 80% ethanol in molecular grade water
    f. Repeat step e for a total of 2 washes.
    g. While the tubes are on the magnet, allow beads to air dry for up to 5 minutes. Note – it is important not to over dry here. Proceed to DNA elution before the beads start to crack.
    h. To elute DNA, remove tubes from the magnetic rack and resuspend beads in Xμl (see below) of 10mM Tris-HCL pH8.0.
    i. Incubate at room temperature for 5 minutes.
    j. Place tubes back on the magnetic rack for 5 minutes and transfer DNA solution to new tube.
75. For downstream qPCR analysis elute in 60-80μl 10mM Tris-HCL pH8.0 and use 1μl per qPCR reaction.
76. For downstream NGS elute in 20μl 10mM Tris-HCL pH8.0.
    a. Use 1μl of eluate to determine the DNA concentration using a Qubit fluorometer.

b. Use 2μl for qPCR analyses of positive and negative control regions (dilute 4x to give final volume of 8μl and use 1μl per reaction).
c. Use remaining eluate (17μl) to prepare libraries using NEBNext® Ultra™ II DNA Library Prep Kit or similar following manufacturer's instructions.
